# Concordance Among Indices of Intrinsic Brain Function: Insights from Inter-Individual Variation and Temporal Dynamics

**DOI:** 10.1101/048405

**Authors:** Chao-Gan Yan, Zhen Yang, Stanley J. Colcombe, Xi-Nian Zuo, Michael P. Milham

## Abstract

Various resting-state fMRI (R-fMRI) measures have been developed to characterize intrinsic brain activity. While each of these measures has gained a growing presence in the literature, questions remain regarding the common and unique aspects these indices capture. The present work provided a comprehensive examination of inter-individual variation and intra-individual temporal variation for commonly used measures, including fractional amplitude of low frequency fluctuations, regional homogeneity, voxel-mirrored homotopic connectivity, network centrality and global signal correlation. Regardless of whether examining intra-individual or inter-individual variation, we found that these definitionally distinct R-fMRI indices tend to exhibit a relatively high degree of covariation, which doesn’t exist in phase randomized surrogate data. As a measure of intrinsic brain function, concordance for R-fMRI indices was negatively correlated with age across individuals (i.e., concordance among functional indices decreased with age). To understand the functional significance of concordance, we noted that higher concordance was generally associated with higher strengths of R-fMRI indices, regardless of whether looking through the lens of inter-individual (i.e., high vs. low concordance participants) or intra-individual (i.e., high vs. low concordance states identified via temporal dynamic analyses) differences. We also noted a linear increase in functional concordance together with the R-fMRI indices through the scan, which may suggest a decrease in arousal. The current study demonstrated an enriched picture regarding the relationship among the R-fMRI indices, as well as provided new insights in examining dynamic states within and between individuals.

## INTRODUCTION

Intrinsic brain activity (IBA) encompasses ongoing neural and metabolic activity and is posited to play a central role in human brain function (Raichle, 2015). In particular, it is believed to help maintain a state of preparedness through the continuous representation of high probability patterns of functional interactions at the local and systems levels, regardless of stimulation level or task demands (Raichle, 2010, 2015). In support of the proposed functional significance of IBA, prior work has linked intra-individual variations in behavior to the state of spontaneous activity in task-positive (e.g., attention) and task-negative (e.g., default) networks (Fox et al., 2007; Kelly et al., 2008). Additionally, a growing literature has worked to link inter-individual variations in trait IBA properties (e.g., functional connectivity) to differences in complex behavioral phenomena (e.g., personality, social reciprocity, intelligence, anxiety) among individuals (see reviews in Di Martino et al., 2014a; Fox and Raichle, 2007; Kelly et al., 2012). With the increased efforts to characterize IBA differences in healthy and clinical populations, a growing number of measures have been proposed – each posited to reveal a distinct aspect of intrinsic brain function (see reviews in Craddock et al., 2013a; Margulies et al., 2010; Zuo and Xing, 2014). While definitional differences exist among these various measures from a theoretical perspective, practical distinctions and commonalities among these measures are yet to be fully explored empirically.

The most fundamental IBA measures are those focused on regional fluctuation properties. Among them, amplitude-based measures have gained particular attention and can be carried out in either the temporal [e.g., resting state fluctuation amplitude (RSFA, Kannurpatti and Biswal, 2008)] or frequency domains [e.g., amplitude of low frequency fluctuations (ALFF, Zang et al., 2007); fractional ALFF (fALFF, Zou et al., 2008); Hurst exponent (Maxim et al., 2005)]. Although commonly used, RSFA and ALFF are somewhat problematic due to their high sensitivity to vascular activity in addition to neuronal signals (Tsvetanov et al., 2015; Vigneau-Roy et al., 2014). Complementing to these regional measures is functional connectivity which indexes the inter-regional synchronization of IBA using measures of temporal correlation or spatiotemporal component decompositions, such as independent component analysis (ICA, Beckmann et al., 2005)]. Building on top of the inter-regional synchronization measures are those focused on particular theoretical constructs, such as local synchronization [e.g., regional homogeneity (ReHo, Zang et al., 2004; Zuo et al., 2013)], interhemispheric synchronization [e.g., voxel-mirrored homotopic connectivity (VMHC, Anderson et al., 2011; Zuo et al., 2010b)] or network centrality [e.g., degree centrality (DC, Buckner et al., 2009; Zuo et al., 2012)]. While each of these measures is widely employed to examine individual and group differences, they are commonly studied in isolation of one another. As a result, questions remain regarding how similar or distinct the various IBA indices are, especially in the absence of an integrative neurophysiological perspective of their underlying mechanisms.

Realizing the potential for differences and overlaps among these indices, some studies have attempted to include a range of measures in their examination of individual-or group-differences. For example, Yan et al. (2013a; 2013b) examined inter-individual differences in ALFF, fALFF, ReHo, DC, and intrinsic functional connectivity (iFC) with a focus on devising strategies for reducing and standardizing their variability. Chen and Xu et al. (2015) systematically investigated various IBA metrics regarding their intra-individual, inter-individual variability and test-retest reliability using richly sampled R-fMRI datasets. In a recent study, fALFF, ReHo and DC were observed to converge at the left angular gyrus with respect to across-individual correlations with working memory maintenance (Yang et al., 2015). Similarly, findings for an array of IBA measures converged in the insula and posterior cingulate cortex in patients with Autism Spectrum Disorders (Di Martino et al., 2014b). Such convergences suggest the presence of a more fundamental mechanism underlying different IBA measures and their variations across individuals.

The goal of the present work is to provide a comprehensive understanding of interdependencies among different IBA measures within and across individuals. In service of this goal, we identified three potential strategies: 1) *Examine how concordant different indices are with respect to variations across voxels.* From this perspective, prior work has found that ALFF is strongly coupled with fALFF (Zou et al., 2008; Zuo et al., 2010a), and also positively correlated with ReHo across voxels (Nugent et al., 2015; Yuan et al., 2013). In a recent study, Aiello et al. (2015) examined the inter-correlation between fALFF, ReHo and DC across voxels, finding high pair-wise correlations among the three measures. 2) *Examine how concordant different indices are with respect to their variation from one individual to the next (i.e., are they capturing unique or shared inter-individual variation?).* Aiello et al. (2015) also provided initial insights in this regard, as they comprehensively examined correlations among fALFF, ReHo and DC across participants, finding high correlations for all gray matter voxels. Beyond that, Yang et al. (2016) used a fully data-driven discovery-replication framework to examine inter-individual differences in six IBA measures (ALFF, fALFF, ReHo, ReHo-2, DC and eigenvector centrality) and six morphology measures, mapping a set of interdependencies of the IBA metrics and their morphological associations. 3) *Examine how concordant different indices are with respect to variation over time (e.g., from one time window to the next within a given individual), a perspective which has yet to be investigated.*

While optimal methodologies are still being determined (Hindriks et al., 2016), the temporal dynamic perspective of IBA measures makes it possible to determine how different measures can be coupled together within an individual. A growing number of studies have demonstrated temporally dynamic changes in iFC patterns (Allen et al., 2014; Chang and Glover, 2010; Handwerker et al., 2012; Hutchison et al., 2013; Kang et al., 2011; Kiviniemi et al., 2011; Smith et al., 2012; Yang et al., 2014). Based on the spatiotemporal structures of iFC, brain activity can be decoded into distinct brain states (Allen et al., 2014; Yang et al., 2014; Zalesky et al., 2014). Dynamic functional network analysis also showed that the dynamic FC fluctuations coincide with periods of high and low network modularity in IBA (Betzel et al., 2016). However, the temporal dynamics of regional IBA as well as their interdependencies remain unexplored.

The present work employed the publicly available Enhanced Nathan Kline Institute-Rockland Sample dataset. Compared to conventional imaging sequences, this fast-sampling multiband dataset is optimal for temporal dynamics analyses given the high temporal resolution (TR = 0.645s). Similar to prior work (Aiello et al., 2015), we first examined the inter-correlation and concordance of differing R-fMRI measures (ALFF/fALFF, ReHo, DC and VMHC) across participants. We then investigated the temporal dynamics of these measures and the extent to which their variations over time are coupled with one another. To determine if a common driving force (either physiological or neuronal) drives their temporal dynamics, we investigated the time varying pattern of global signal correlation (GSCorr, Power et al., 2016). We also tested for any potential relationships with physiological noises (respirational volume and heart rate). Finally, to examine their potential value in neuronal applications, we investigated how coupling relationships change with age across the lifespan

## MATERIALS AND METHODS

### Participants and Imaging Protocols

Multiband fast-sampling R-fMRI data (TR = 0.645s, 9.675 minutes/900 volumes) of 173 neurotypical individuals between ages 8.0 and 86.0 (mean age: 44.5 years; 117 females) with datasets passing quality criteria were selected from the publicly available Enhanced Nathan Kline Institute-Rockland Sample (Available via International Neuroimaging Data-sharing Initiative: http://fcon_1000.projects.nitrc.org/indi/enhanced/). Images were acquired in a 3T Siemens TIM Trio scanner using the following multiband echo-planar imaging (EPI) sequence: TE = 30 ms, flip angle = 60°, slice thickness = 3.0 mm, field of view = 222mm, matrix size = 74×74, TR = 645ms; no gap, resolution = 3.0×3.0×3.0 mm^3^. During data acquisition, participants were instructed to simply rest with eyes open (Nooner et al., 2012). For spatial normalization and localization, a high-resolution T1-weighted magnetization prepared gradient echo image (MPRAGE) was also obtained for each participant. To monitor physiological signals, cardiac signal was recorded using a pulse oximeter (BIOPAC) placed on the right index finger. Respiration was recorded using a respiratory transducer (BIOPAC) with belt placed around the abdomen. These physiological data were sampled every 16 ms during fMRI scanning. Of the 173 participants, 22 were missing both respiratory and cardiac recordings, while 39 were missing either cardiac or respiratory recordings. The Nathan Kline Institute institutional review board approved the submission of anonymized data obtained with written informed consent from each participant.

### Preprocessing

Unless otherwise stated, all preprocessing was performed using the Data Processing Assistant for Resting-State fMRI (DPARSF, Yan and Zang, 2010, http://rfmri.org/DPARSF), which is based on Statistical Parametric Mapping (SPM8) (http://www.fil.ion.ucl.ac.uk/spm) and the toolbox for Data Processing & Analysis of Brain Imaging (DPABI, Yan et al., 2016, http://rfmri.org/DPABI). The first 16 volumes (10.32s) were removed to allow data to reach equilibrium, leaving a total of 884 volumes for final analysis. The functional volumes for each subject were motion-corrected using a six-parameter (rigid body) linear transformation with a two-pass procedure (registered to the first image and then registered to the mean of the images after the first realignment). Individual structural images (T1-weighted MPRAGE) were co-registered to the mean functional image after realignment using a 6 degrees-of-freedom linear transformation without re-sampling. The transformed structural images were then segmented into gray matter (GM), white matter (WM) and cerebrospinal fluid (CSF) (Ashburner and Friston, 2005). Based on these segmented images, the Diffeomorphic Anatomical Registration Through Exponentiated Lie algebra (DARTEL) tool (Ashburner, 2007) was used to compute transformations from individual native space to MNI space. Gray matter density (GMD) maps were also used as covariates in group analyses to control morphometric contributions to age effects.

### Nuisance Regression

Recent work has demonstrated that micro head movements, as small as 0.1mm between time points, can introduce systematic artifactual inter-individual and inter-group variability in R-fMRI measures (Power et al., 2012a, b; Satterthwaite et al., 2012; Van Dijk et al., 2012). Here we only selected participants with relatively low head motion, as assessed using mean frame-wise displacement (derived from Jenkinson’s formula, Jenkinson et al., 2002) (criteria: mean FD < 0.2mm). We also utilized the Friston 24-parameter model (Friston et al., 1996) to regress out head motion effects from the realigned data (i.e., 6 head motion parameters, 6 head motion parameters one time point before, and the 12 corresponding squared items) based on recent reports that higher-order models demonstrate benefits in removing head motion effects (Satterthwaite et al., 2013; Yan et al., 2013a). Head motion was also controlled at the group-level by taking mean FD as a covariate (Satterthwaite et al., 2013; Yan et al., 2013a).

The signals from WM and CSF were regressed out to reduce respiratory and cardiac effects. In addition, linear and quadratic trends were also included as regressors since the blood oxygen level dependent (BOLD) signal demonstrates low-frequency drifts. Global signal regression (GSR) was not performed because of concerns about increasing negative correlations (Murphy et al., 2009; Weissenbacher et al., 2009) and possible distortions in group differences in iFC (Gotts et al., 2013; Saad et al., 2012). Temporal filtering was not performed to preserve the temporal dynamics of time series, but linear detrend was performed. After nuisance regression, the functional data were registered into MNI152 space with 3mm^3^ cubic voxels by using the transformation information acquired from the previous DARTEL step.

### R-fMRI Indices for Regional Characteristics and Functional Synchrony

We examined the interdependency among the following five R-fMRI based indices of intrinsic brain activity:

1) Amplitude measures: ALFF (Zang et al., 2007) and fALFF (Zou et al., 2008). ALFF is the mean of amplitudes within a specific low frequency range (0.01-0.1 Hz in the current study) from a Fourier decomposition of the time course. fALFF is the ratio of the sum of amplitudes of a given low frequency band (here, 0.01-0.1Hz) to the sum of Fourier amplitudes across the entire frequency range. ALFF is proportional to the strength or intensity of low frequency oscillations, while fALFF represents the relative contribution of specific oscillations to the whole detectable frequency range.

2) Regional homogeneity (ReHo, Zang et al., 2004). ReHo assesses the degree of regional synchronization/coherence among fMRI time courses. It is defined as the Kendall’s coefficient of concordance (or Kendall’s W, Kendall and Gibbons, 1990) between the time series of a given voxel with those of its nearest neighbors (26 in the current study).

3) Voxel-mirrored homotopic connectivity (VMHC, Anderson et al., 2011; Zuo et al., 2010b). VMHC corresponds to the functional connectivity between any pair of symmetric inter-hemispheric voxels-that is, the Pearson’s correlation coefficient between the time series of each voxel and that of its counterpart voxel at the same location in the opposite hemisphere. The resultant VMHC values were Fisher-Z transformed. For better correspondence between symmetric voxels, VMHC requires that individual functional data are first registered in MNI space and smoothed (4.5 mm FWHM) and then registered to a symmetric template. The group averaged symmetric template was created by first computing a mean normalized T1 image across participants, and then this image was averaged with its left–right mirrored version (Zuo et al., 2010b).

4) Network degree centrality (Buckner et al., 2009; Zuo et al., 2012). Degree centrality is the number or sum of weights of significant connections for a voxel. Here, we generated voxel-based whole-brain correlation for a given voxel, and then calculated the weighted sum of positive correlations by requiring each connection’s correlation coefficient to exceed a threshold of r > 0.25 (p < 0.05, Buckner et al., 2009).

5) Global Signal correlation (GSCorr): GSCorr is the Pearson correlation coefficient between the global mean time series (averaged across all voxels inside the group mask^1^) and each voxel in the group mask. These correlation values were then Fisher-Z transformed.

### Dynamic R-fMRI Indices

Dynamic R-fMRI indices were generated using sliding time-window analysis. First, we applied hamming windows (length of 100 TRs, overlapping by 3 TRs)^2^ to BOLD signals to obtain windowed time series; second, within each window, we calculated the above-mentioned R-fMRI indices (i.e., fALFF, ReHo, VMHC, DC and GSCorr). To characterize the dynamic R-fMRI indices, we computed the mean and standard deviation (SD) map across time windows for each index. To quantify temporal variations of these indices, we also computed the coefficient of variation (CV: SD/mean) map over time for each index. The CV maps were then Z-standardized relative to their own mean and SD across all voxels within the group mask. Across participants, we performed one-sample t-tests on the Z maps of CV to investigate the overall pattern of temporal variation.

### Validation of Temporal Dynamics Through Simulation

A few recent studies have raised concerns regarding the study of temporal dynamics using R-fMRI data due to the potential for artifactual findings (Hindriks et al., 2016; Laumann et al., 2016). In particular, Hindriks et al. noted that observed temporal variation in an estimated statistic does not imply that the underlying population value is variable (Hindriks et al., 2016); this work suggested the need to raise the bar for temporal dynamics data analysis by using appropriately randomized data to establish the contributions of noise. Here, to address this concern, we created surrogate data by randomizing the Fourier phases but keeping the amplitude of the real time-series data, and then performed the same analysis to confirm that our findings exist on surrogate data.

### Correlation between Global Mean of R-fMRI Indices

For static analysis, the global mean of each index was computed within the group GM mask^3^ for each of the 173 participants. Pearson’s correlation coefficient was then computed between pairs of the global mean of R-fMRI indices across participants. For dynamic analysis, we calculated the global mean of a given R-fMRI measure for each time window. This resulted in one global mean time series per index per participant. Within each participant, we calculated the correlations between pairs of the global mean time series of R-fMRI indices. These correlations were then averaged across participants to provide an estimation of the overall dynamic correlations. To examine if the temporal dynamics of these measures were being driven by physiological noise, we examined their relationship with each: motion, global signal, respiration volume per time (RVT) and heart rate (HR). RVT and HR were extracted by applying AFNI’s RetroTS program to the cardiac and respiratory signals. RVT/HR was averaged within each time window before usage in correlation analyses carried out across windows.

### Voxel-wise Concordance Index

We calculated the Kendall’s W of the five static R-fMRI indices (with 4.5 mm smoothing^4^) across participants for each voxel as a static voxel-wise concordance index. Kendall’s W was used because this non-parametric statistic has no assumptions of the distribution and is insensitive to differences in scale among the R-fMRI indices. Given the high colinearity between fALFF and ALFF, we chose to include only one of the two measures in the concordance indices to avoid artificially inflating the concordance measure. fALFF was chosen because it is less susceptible to artifactual contributions of motion and pulsatile effects as compared with ALFF (Yan et al., 2013a; Zou et al., 2008; Zuo et al., 2010a). To compare the within-individual dynamic concordance to the static functional concordance across individuals, we calculated Kendall’s W for the five R-fMRI indices across time windows as the dynamic voxel-wise concordance index. A one-sample t-test was performed on the Z-standardized dynamic voxel-wise concordance index (with 4.5 mm smoothing^5^) to examine the pattern among subjects. The rank order of the static voxel-wise concordance as well as the rank order of the group t map of dynamic voxel-wise concordance index were calculated to compare their consistency.

### Volume-wise Concordance Index

For static analyses, we calculated Kendall’s W of the R-fMRI indices across all brain voxels for each participant. To investigate the properties of functional concordance, we compared subjects with the highest 20% static volume-wise concordance to those with the lowest 20% static volume-wise concordance. For each R-fMRI measure, two-sample t-tests were performed between the highest 20% and lowest 20% subjects to capture group differences. For dynamic analyses, a dynamic volume-wise concordance index was computed the same way as static analysis, but for each time window. The mean dynamic spatial functional coupling index (across time) was then compared with the static volume-wise concordance index. To characterize the properties of high concordance states in contrast with low concordance states at the within-individual level, we compared properties of the top 20% windows with the highest dynamic volume-wise concordance to those windows with the lowest 20% dynamic volume-wise concordance. For each individual, a mean map of a given measure was calculated for the top 20% windows, as well as a mean map for the lowest 20% windows. Then paired t-tests were performed to compare differences in properties between high windows and low windows.

### Age Effects on Functional Concordance

Finally, to examine the potential role of functional concordance in development, we linked the functional concordance indices to age across lifespan. A general linear model with sex and motion as covariates was utilized within DPABI. Specifically, for voxel-wise or volume-wise concordance indices, the following regression model was constructed:

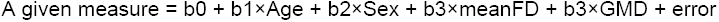

Mean FD was included to control for the residual effect of head motion. We opted to employ group-level corrections over scrubbing approaches for the following reasons: (1) recent work suggested that scrubbing offers little advantage over group-level motion correction combined with Friston 24 model at the individual level. Scrubbing can also be less conservative when motion correlates with a between-subject variable of interest, and can be incomplete in their corrective effects (Yan et al., 2013a); (2) scrubbing of non-contiguous time points alters the underlying temporal structure of the data, which can unintentionally bias estimates of an individual’s intrinsic activity patterns towards one state over another in temporal dynamics studies. In addition to sex and head motion, we also included gray matter density (GMD) as a covariate to control for the effects of structural differences on the correlation between age and functional concordance. In response to recent concerns regarding cluster-based multiple comparison corrections (Eklund et al., 2016), we utilized permutation test (permuted 1000 times) with Threshold-Free Cluster Enhancement (TFCE, Smith and Nichols, 2009) to threshold our results at p < 0.05 corrected. TFCE is a strict multiple comparison correction strategy implemented in PALM (Winkler et al., 2016) and integrated into DPABI (Yan et al., 2016).

## RESULTS

### Static and Dynamic R-fMRI Indices

The first step in the present work was to characterize the temporal variation in our R-fMRI indices of interest. In this regard, for each of the indices, we carried out sliding window analyses (window size: 100 TRs = 64.5 s) and then calculated the voxel-wise coefficient of variation across windows for each participant. The resultant coefficient of variation maps are depicted in Figure 1, along with traditional static R-fMRI index maps. The CV maps revealed different patterns of variation for different R-fMRI indices. ALFF/fALFF appeared to show lower temporal variation in high level cognitive regions (e.g., default mode network components, dorsolateral prefrontal cortex), and higher variation in lower-level primary sensory and motor areas, as well as subcortical regions. In contrast, for ReHo and DC, the variation is high among all the cortices, including the default mode network. Of note, CV is not applicable for VMHC and GSCorr as they included negative correlation values.

**Figure 1.**
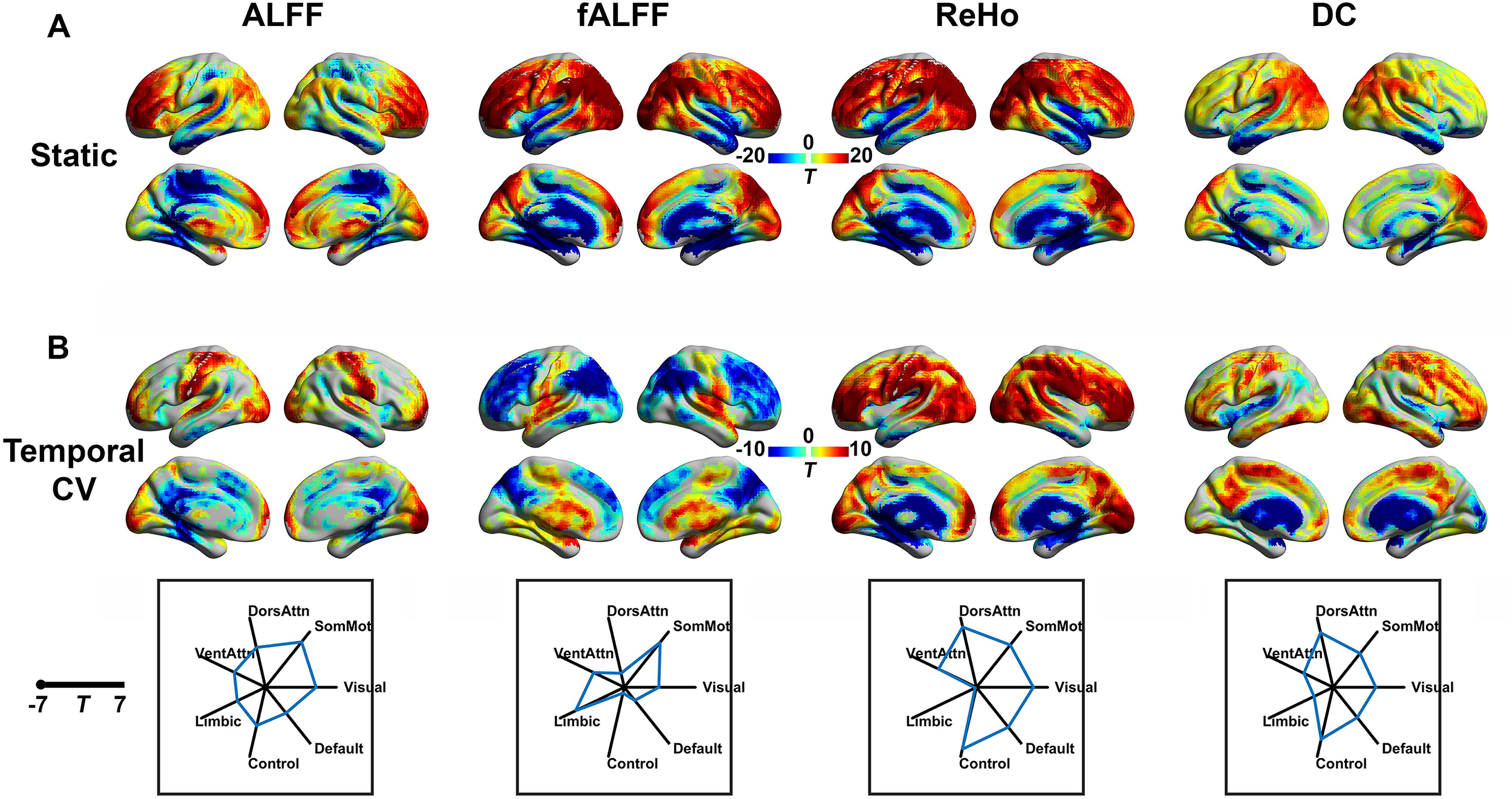
The spatial maps for static and dynamic R-fMRI indices. (A) T maps of static indices across 173 subjects, which were corrected using permutation test (permuted 1000 times) with threshold-free cluster enhancement (TFCE) at p < 0.05. (B) T maps of temporal coefficients of variation (CV) of dynamic R-fMRI indices, which were corrected using the same permutation with TFCE at p < 0.05. In the bottom row, t values of Z-standardized temporal CV of dynamic R-fMRI indices were averaged within each of the seven networks defined by Yeo et al. (2011) and plotted using radar plots. ALFF: amplitude of low frequency fluctuations; fALFF: fractional ALFF; ReHo: regional homogeneity; DC: degree centrality; SomMot: somatomotor; DorsAttn: dorsal attention; VentAttn: ventral attention; Control: frontoparietal control; the abbreviations for R-fMRI indices and seven networks will be used in the same way in all following figures. Of note, CV is not applicable for VMHC (voxel-mirrored homotopic connectivity) and GSCorr (global signal correlation) as they included negative correlation values, thus they were not plotted.

### Evaluating Concordance among R-fMRI Indices: Global-Level Analyses

From the perspective of inter-individual variation, all measures were closely coupled with each other at the global level. Static analyses, focused on the mean value within the grey matter mask for each of the measures of interest at the individual level, found high correlation between each pair of R-fMRI indices across individuals (Figure 2A). Dynamic analyses capitalized on the ability to perform within-individual analyses across windows; we found a high degree of correlation between all pairs of R-fMRI indices over time (mean correlation scores across 112 participants who had both cardiac and respiratory recordings are depicted in Figure 2B), indicating common fluctuation patterns underlying all the functional aspects. Of note, GSCorr is less correlated with ALFF and fALFF (mean r=0.62 and 0.58), but highly correlated with DC (mean r = 0.88). Nuisance parameters, including motion, RVT and HR did not appear to be major driving forces for the observed ensemble changes. The temporal changes did not correlate with the global signal itself (Figure 2B).

**Figure 2.**
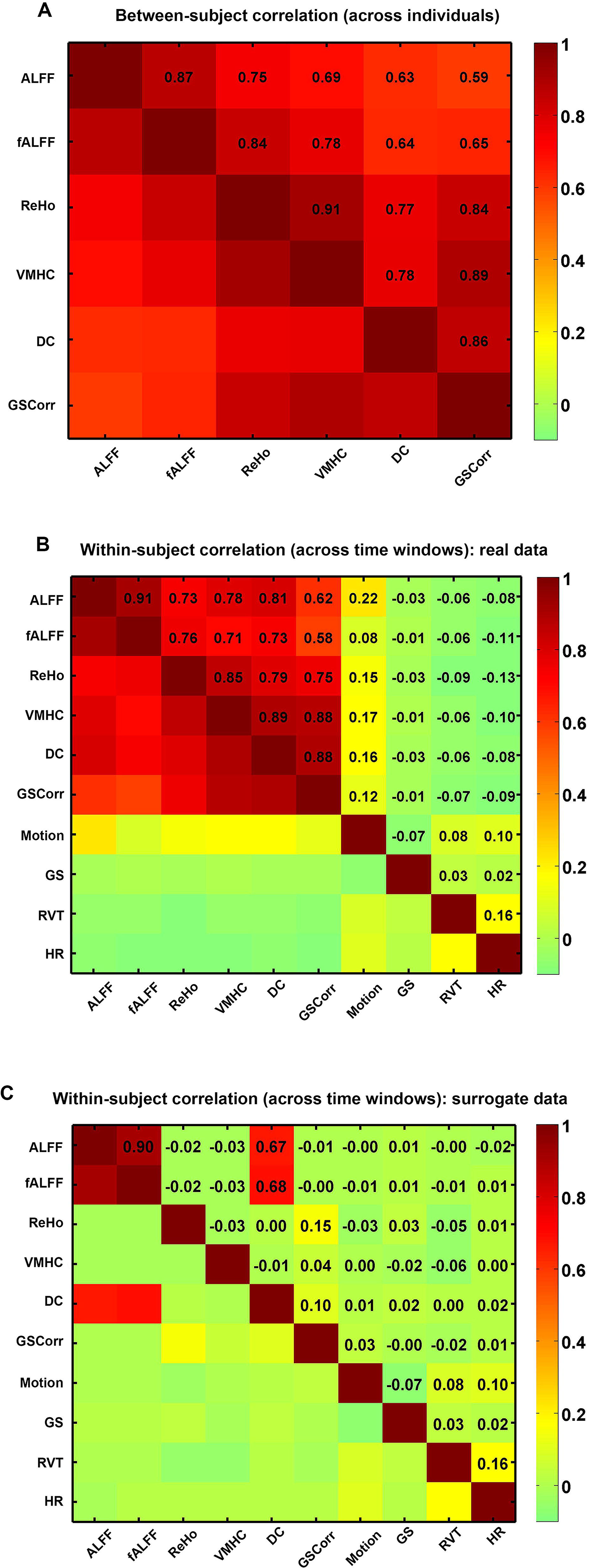
Global-level concordance among R-fMRI indices. (A) Static concordance: Pearson’s correlations between pairs of global means of static R-fMRI indices computed across participants. (B) Dynamic concordance on real data: Pearson’s correlations computed between pairs of global mean time series derived from dynamic R-fMRI indices within a participant and then averaged across participants. Their association with nuisance variables were also examined: motion, global signal (GS), respiration volume per time (RVT) and heart rate (HR). (C) Dynamic concordance on surrogate data (phase randomized). Pearson’s correlations similarly computed as in (B) except for on phase randomized surrogate data.

Next, we worked to verify that the high correlations observed at the global level between pairs of R-fMRI indices over time were not caused by random variations (Hindriks et al., 2016). To accomplish this, we performed the same temporal dynamical analyses on surrogate data, which was created by randomizing the Fourier phases of the time-series data. As demonstrated in Figure 2C, ALFF and fALFF had high correlation (r = 0.9) on surrogate data – this was to be expected given their definitional similarity. Such a correlation doesn’t reflect any neuronal synchronization given the similarity in deriving the measures of ALFF and fALFF. Additionally, DC has relatively high correlation with ALFF and fALFF on surrogate data – a finding consistent with the work of Tomasi et al. (2016). No prominent correlations were observed between any of the other pairings of measures applied to surrogate data – this suggests that the common fluctuation patterns underlying the various functional indices are not driven by nuisance signal or random variations.

### Evaluating Concordance among R-fMRI Indices: Voxel-wise Analyses

From a static perspective, at each voxel, we calculated the Kendall’s W (coefficient of concordance) of the five R-fMRI indices (fALFF/ReHo/VMHC/DC/GSCorr) across participants. Of note, we did not include ALFF here to avoid artificially inflating the concordance measure, given its extremely high correlation with fALFF. As demonstrated in the concordance map (Figure 3A), gray matter regions exhibited high concordance across individuals, including cerebral cortex and subcortical regions. In contrast, white matter regions demonstrated low concordance. From a temporal dynamic perspective, we calculated the voxel-wise Kendall’s W of the five R-fMRI indices across time windows within each individual. As demonstrated in the one-sample t-test on the Z-standardized within-individual concordance maps (Figure 3B), these analyses revealed that the voxel-wise concordance maps are highly similar to the static concordance. The rank order of the static concordance (Figure 3C) and the rank order of the dynamic concordance (Figure 3D) were highly correlated (r = 0.84, Figure 3E).

**Figure 3.**
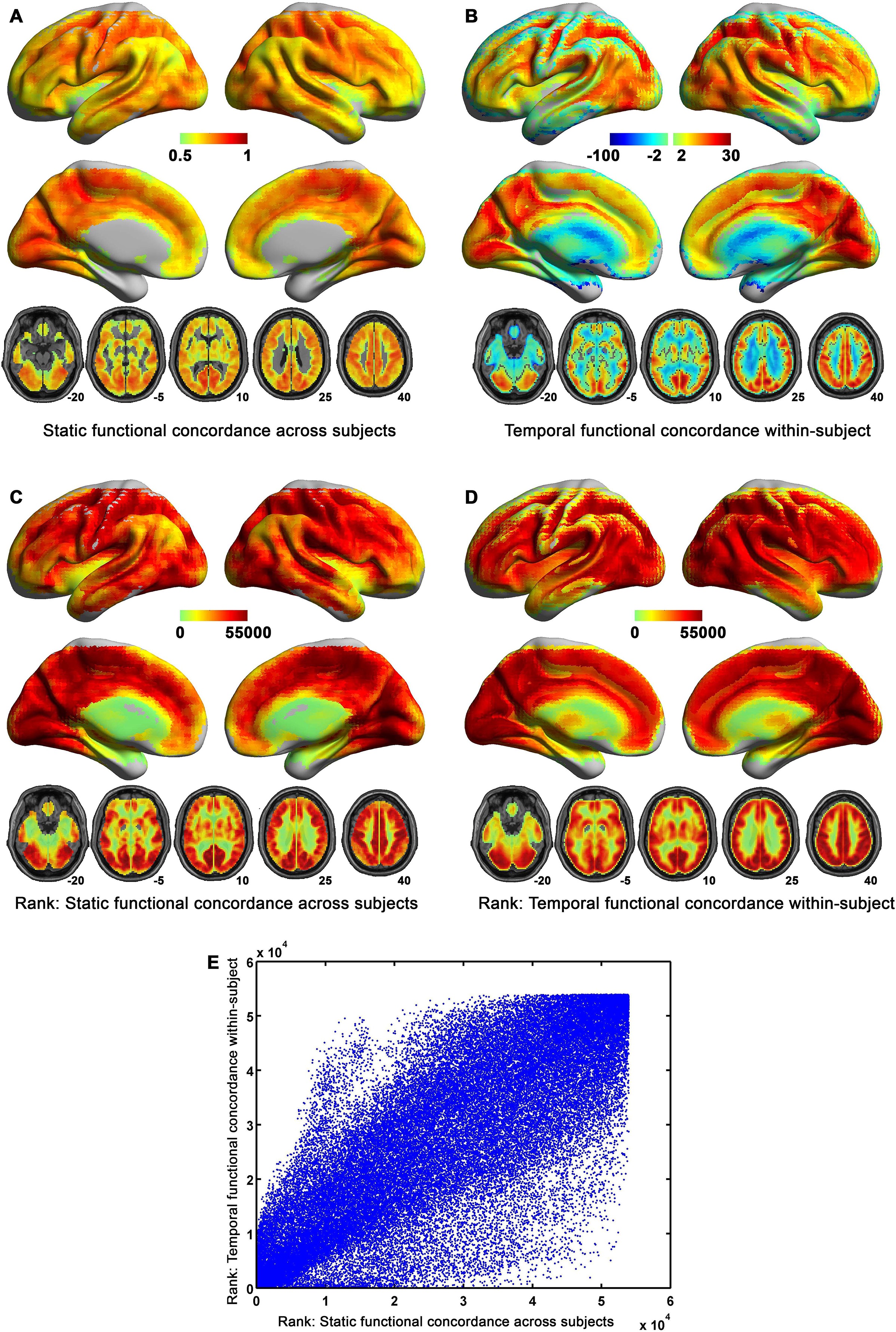
Functional concordance among five R-fMRI indices: voxel-wise analyses. (A) Static concordance among R-fMRI indices computed across participants (Kendall’s W); (B) Dynamic concordance among R-fMRI indices (T scores) computed in two steps: first, in each participant, Kendall’s W was computed across time windows among five R-fMRI indices; then one-sample T-tests were performed to collapse the concordance maps (after Z-standardization) across participants; (C) Rank order of A; (D) Rank order of B; (E) Spatial correlation between C and D.

Importantly, voxel-wise concordance was related to age. As indicated by age effects on the concordance map, the brain showed widespread reductions in functional concordance as age increased, especially in subcortical areas (Figure 4A). This reduction was independent of gray matter density, sex and motion as all those were controlled as covariates. To quantify the overall functional concordance, we averaged the dynamic voxel-wise concordance across gray matter voxels, and found this mean functional concordance index was negatively correlated with age across participants (partial correlation =-0.36, p = 1×10^-6^, controlling for global GMD, sex and motion; Figure 4B).

**Figure 4.**
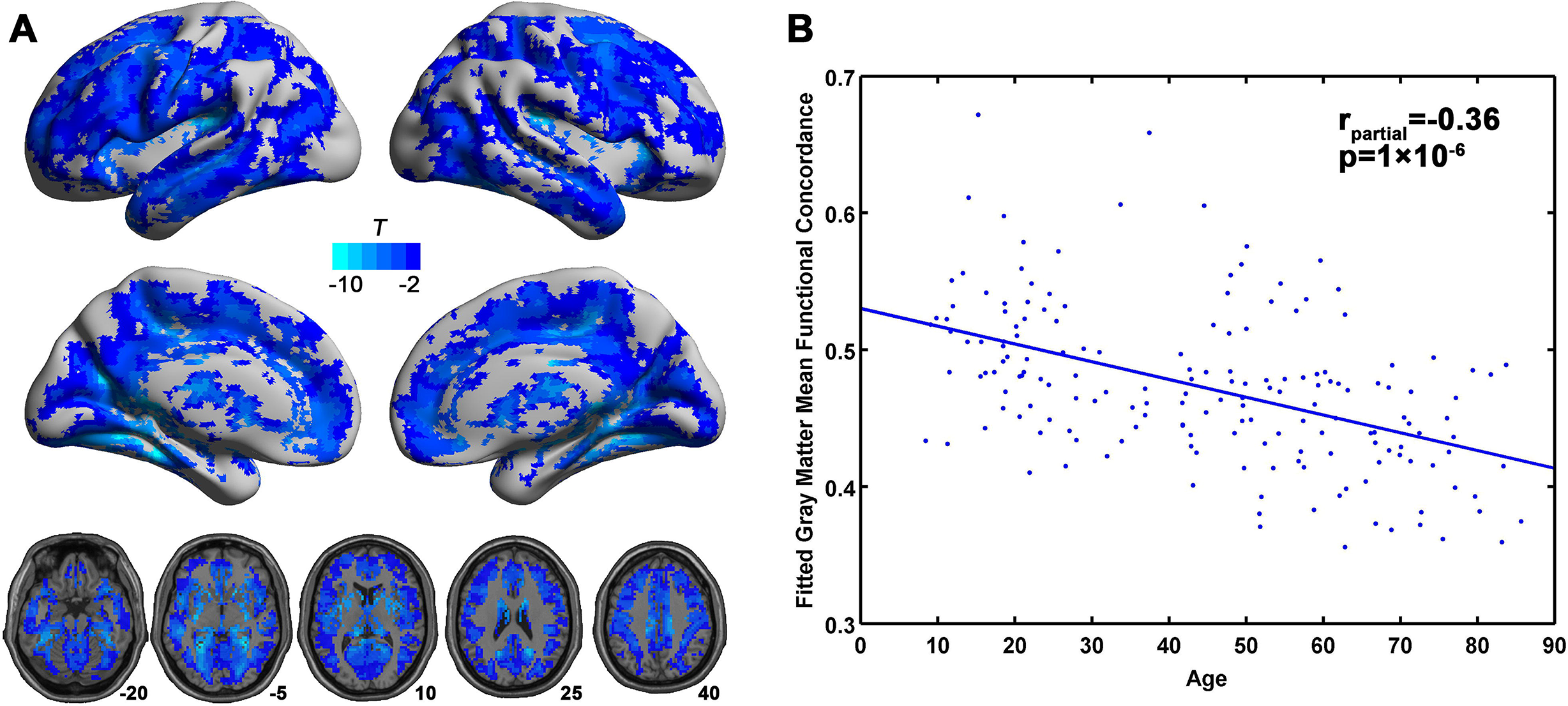
Age effects on dynamic voxel-wise within-individual concordance (Kendall’s W). (A) Widespread reductions in functional concordance noted with increasing age. Results were corrected using permutation test (permuted 1000 times) with threshold-free cluster enhancement (TFCE) at p < 0.05. (B) Mean functional concordance index (dynamic voxel-wise concordance averaged across gray matter voxels) is negatively correlated with age across subjects.

### Evaluating Spatial Concordance among R-fMRI Indices: Volume-wise Analysis

Statically, we calculated the concordance (Kendall’s W) of the five measures (fALFF/ReHo/VMHC/DC/GSCorr) across voxels for each individual, finding that volume-wise static concordance varied across participants, and was negatively correlated with age (partial correlation=-0.42, controlling for global GMD, sex and motion; Figure 5A). Dynamically, the same concordance among the five R-fMRI indices was computed for each time window, yielding a concordance time-series for each participant (see Figure 5B for an example). The CV of dynamic volume-wise concordance was not correlated with age (partial correlation=-0.04, controlling for global GMD, sex and motion; Figure 5C).

**Figure 5.**
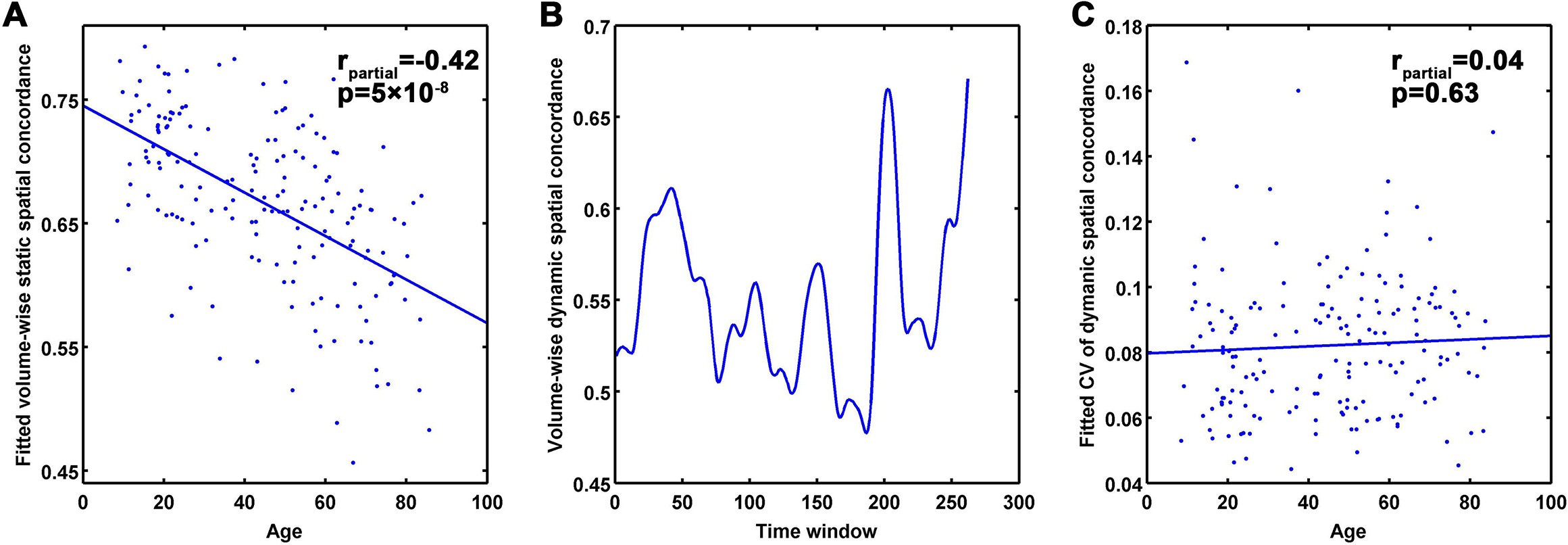
Functional concordance among five R-fMRI Indices: volume-wise analyses. (A) Static volume-wise spatial concordance among R-fMRI indices was plotted as a function of age; (B) A dynamic volume-wise concordance time series for an exemplar subject; (C) the CV of dynamic volume-wise spatial concordance plotted as a function of age.

### Understanding Variations in Concordance

To understand the functional significance of concordance among the different indices, we first divided participants into five groups based on volume-wise static concordance, and then compared the highest and lowest groups using a two-sample t-test. Participants with the highest 20% volume-wise static concordance demonstrated much higher values of all five R-fMRI indices than those with the lowest 20% concordance (Figure 6A)^6^. We tested if head motion could be a primary driving factor for this difference, finding the two groups did not differ significantly in head motion (t =-1.5, p = 0.13).

**Figure 6.**
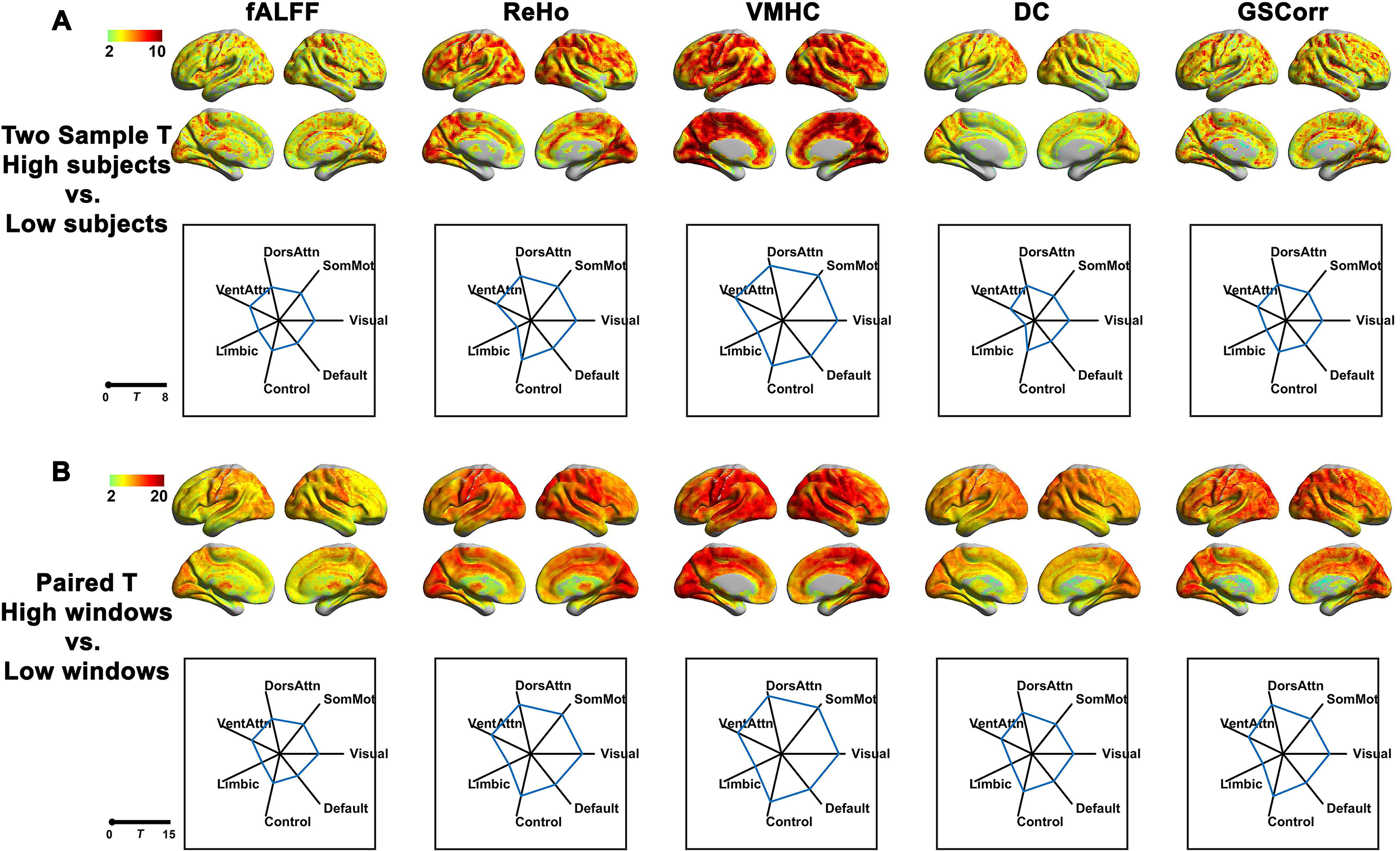
Differences in magnitude of R-fMRI indices between high and low concordance participants or time windows (between the upper 20% and the lower 20%). (A) T-scores obtained by comparing static R-fMRI indices between participants with highest and lowest 20% of concordance (static volume-wise spatial concordance) were plotted for each voxel on surface maps (top row). The average t-scores of each of the seven Yeo et al., (2011) networks were plotted in the radar charts (bottom row). (B) Paired sample t-scores obtained by comparing windows with highest 20% and lowest 20% concordance (dynamic volume-wise spatial concordance) were plotted for each voxel on surface maps (top row). Specifically, for each participant, the mean R-fMRI index maps for the windows with highest 20% and lowest 20% concordance were calculated, yielding two maps (high concordance, low concordance), and then paired t-tests were performed between high and low concordance maps across participants. The average t-score of each network were plotted in radar charts (bottom row). Results were corrected by permutation test (permuted 1000 times) with TFCE at p < 0.05 corrected.

To further investigate relationships between concordance and the strength of R-fMRI indices, we carried out temporal dynamic analyses, which differentiated high and low concordance states on a within-individual basis. Specifically, for each participant, we sorted windows based on their concordance score and we calculated the mean map for the highest 20% and lowest 20% of windows. For each R-fMRI index, this yielded two maps for each participant (high concordance, low concordance). Paired t-tests were employed to identify systematic changes in R-fMRI indices across the two concordance states. The windows with high concordance demonstrated higher scores for all functional aspects (fALFF/ReHo/VMHC/DC/GSCorr) relative to those observed with low concordance (Figure 6B), suggesting high and low integration states exist in the brain. When each of the 7 large-scale networks previously defined by Yeo et al. were examined, the limbic network emerged as a notable exception, showing only modest increases in R-fMRI indices from low to high-integration states. These differences were not driven by random variations, as no significance differences were found in any of the R-fMRI indices between high and low concordance states on phase randomized surrogate data. Of note, as demonstrated by Cohen’s f^2^ in Figure S2, the effect size of findings obtained for the contrast between high and low concordance within individuals (high time windows vs. low time windows) was greater than that between individuals (high subjects vs. low subjects).

Given the seemingly broad nature of increases in connectivity during high concordance states, we further asked whether increases in connectivity were specific to within-network connectivity, or extended between-network connectivity as well. To answer this question, we investigated intra-network functional connectivity and inter-network functional connectivity using the 7 Yeo networks (Yeo et al., 2011). For a pre-defined network, we calculated the mean time course for that network, and then correlated with all the voxels within another network. These correlations were averaged to represent the general correlation between the two networks. The correlation between the mean time course and all voxels within the same network can be averaged to represent the within-network functional connectivity. In general, within network functional connectivity is higher than between-network connectivity (Figure 7A and B). For between-network connectivity, the limbic network demonstrated the lowest connectivity with others. The observation that both intra- and inter-network correlations were higher in the highest 20% windows than the lowest 20% windows (Figure 7C), suggests that both within- and between-network connectivity were increased during higher concordance states.

**Figure 7.**
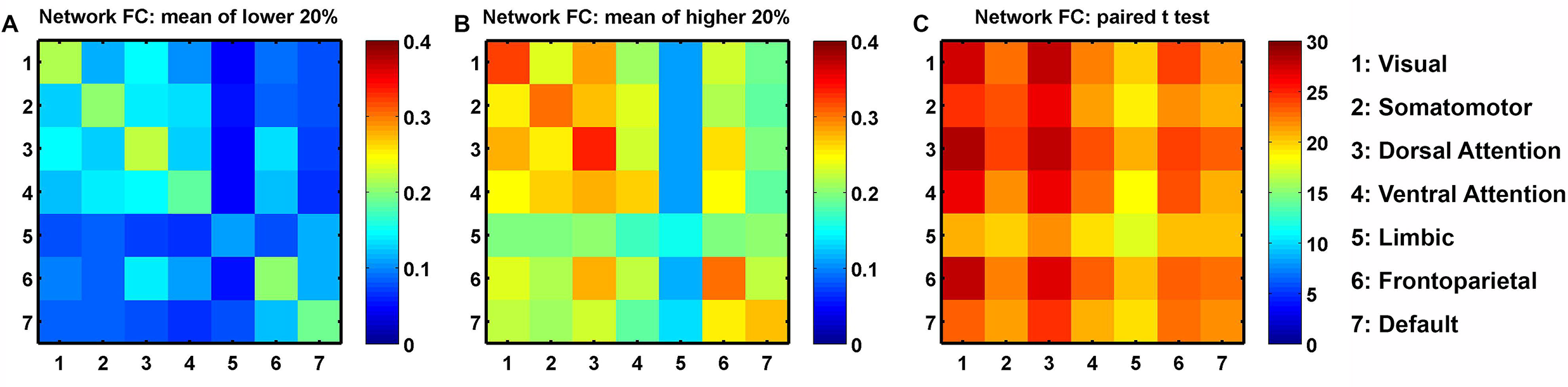
Intra- and inter-network functional connectivity between pairs of Yeo et al., (2011) networks at low (A) and high (B) concordance states. Low and high concordance states were defined based on 20% of windows with the lowest or highest dynamic volume-wise spatial concordance. (C) Paired t-tests of functional connectivity between the 20% highest and lowest windows.

To further address what impacts the fluctuation of concordance, we calculated the correlation between concordance time series and the time series of each R-fMRI measure/physiological signal (Figure 8). The mean correlation (averaged across 112 participants who had both cardiac and respiratory recordings) between concordance and R-fMRI measures were generally high, while the correlation between concordance and head motion (r = 0.047), RVT (r =-0.082) and HR (r =-0.118), were low. These findings confirm that these physiological signals do not appear to be the primary driving force for the observed fluctuations in functional concordance. We also tested if concordance had any temporal associations across time windows. Surprisingly, we found that while high and low concordance windows were present throughout a scan session, the frequency of high concordance windows did appear to increase over time as an individual remained in the scanner (Figure 9A) – even linear detrend has been performed for each voxel before the dynamic R-fMRI indices were performed. The actual increase in concordance was small (mean concordance: 0.56 → 0.59, Figure 9B); nonetheless, they were significant (on average, linear trends explained 15.9% of temporal changes in concordance within participants) and detectable in multiple other datasets that we checked (e.g., Human Connectome Project). We also noted that linear increases exist in all functional aspects we examined (ALFF, fALFF, ReHo, VMHC, DC an GSCorr, Figure 9C~H), which is consistent with an prior study reporting similar increases in low frequency power in a single scanning session (Duff et al., 2008). Importantly, such an increase doesn’t exist in phase randomized surrogate data (Figure S3), suggesting the linear increase might reflect neuronal contributions. Prior studies found increase in global signal amplitude corresponds to decrease in EEG vigilance (Wong et al., 2016; Wong et al., 2013), thus the linear increases in functional concordance as well as other R-fMRI indices in the current study may reflect changes in the intrinsic architecture associated with increase in fatigue or decrease in arousal.

**Figure 8.**
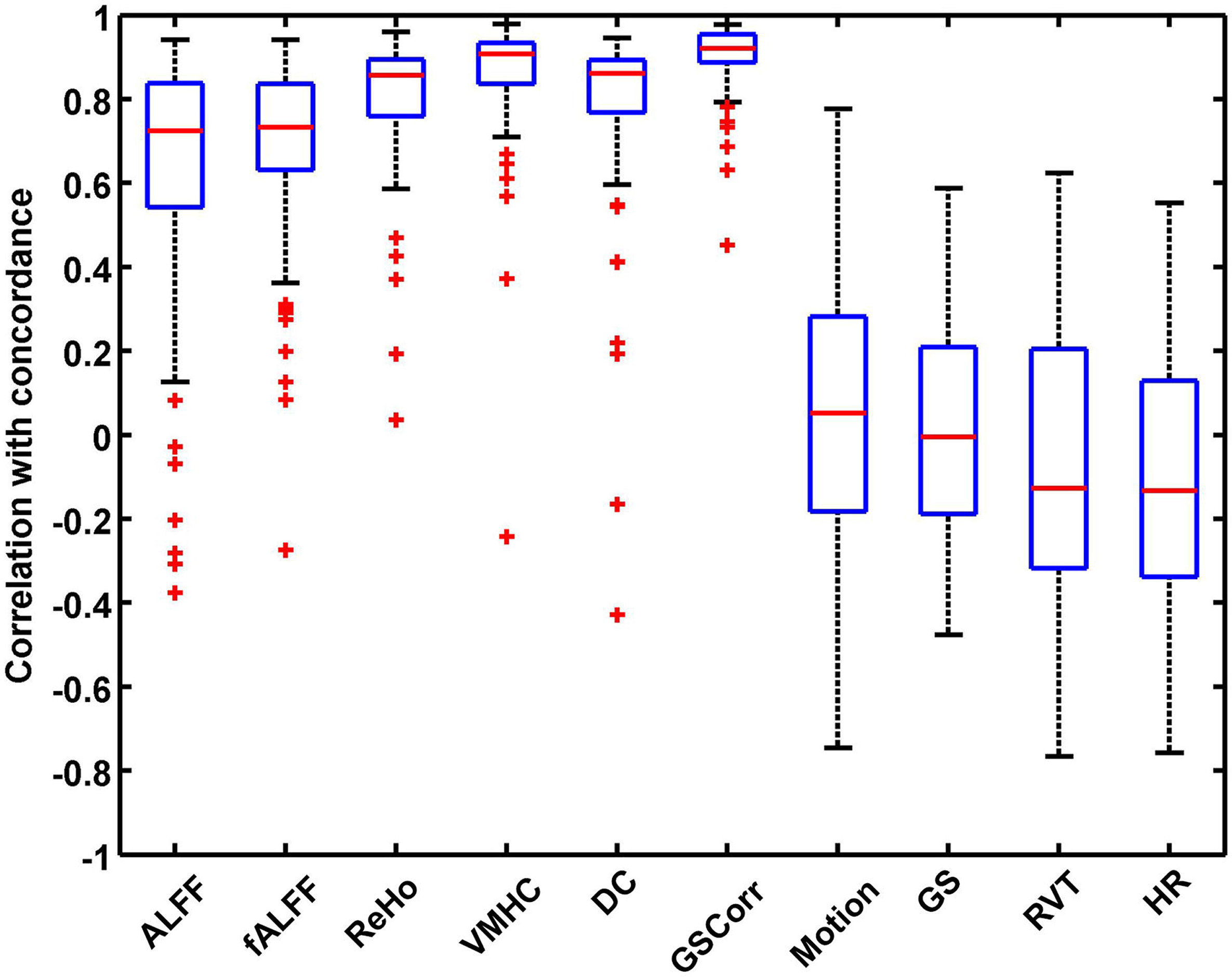
Correlations between concordance time series and time series of each R-fMRI measure or physiological signal. The mean correlation (r; averaged across 112 participants who had both cardiac and respiratory recordings) between concordance and ALFF: 0.643; fALFF: 0.693; ReHo: 0.808; VMHC: 0.865; DC: 0.793; GSCorr: 0.904; Motion: 0.047; GS: 0.005; RVT:-0.082; HR:-0.118. GS: global signal; RVT: respiration volume per time; HR: heart rate.

**Figure 9.**
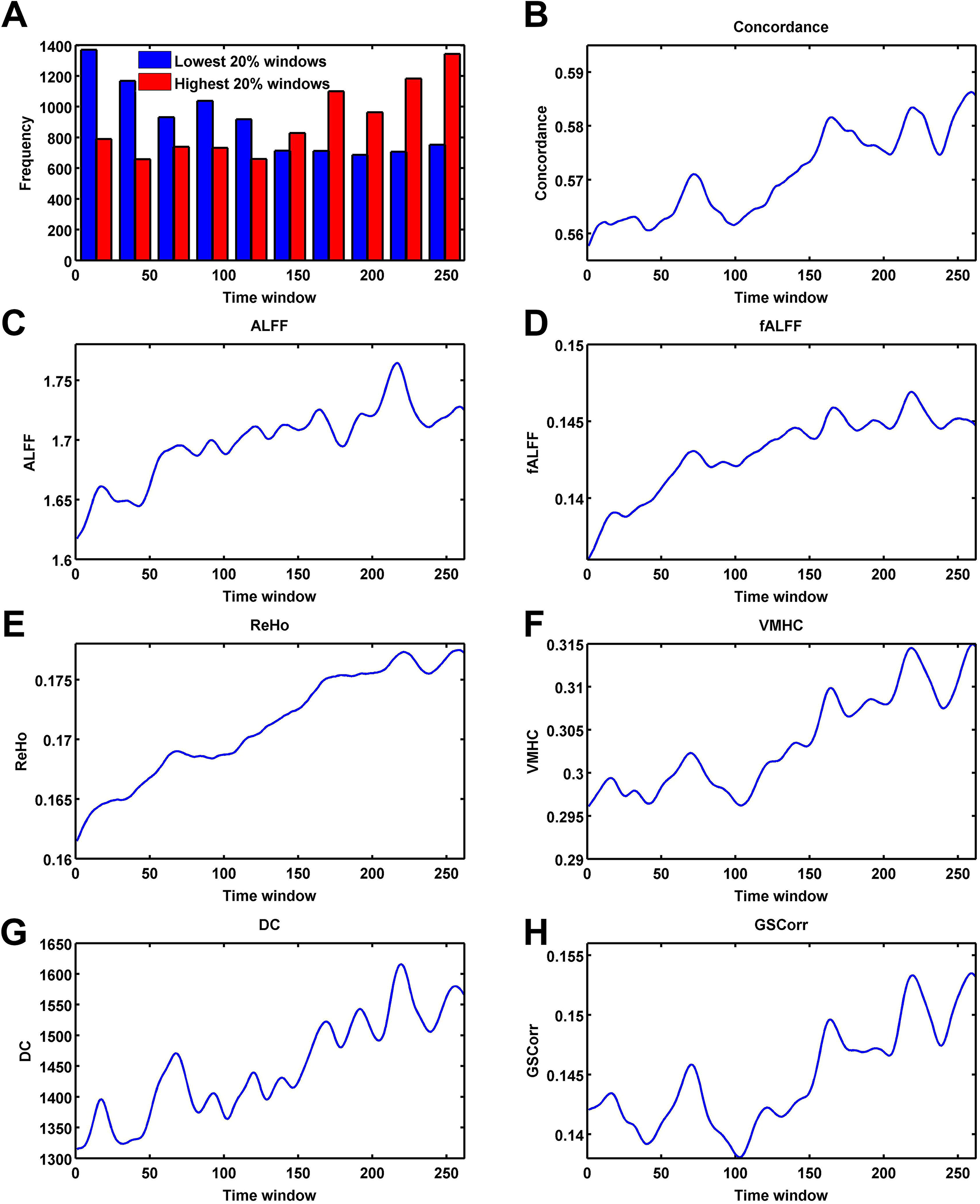
The impact of scanning temporal order on functional concordance. (A) The frequency distribution of time window for the lowest 20% windows and highest 20% windows across 173 participants. The lowest 20% windows occurred more frequently at the beginning of the scanning, while the highest 20% windows occurred more frequently at the end. (B) Mean concordance time course averaged across 173 participants. The concordance increased significantly over time. Linear trend also exists in all R-fMRI indices: mean time course of ALFF, fALFF, ReHo, VMHC, DC and GSCorr averaged across 173 participants (C-H).

## DISCUSSION

The present work provided a comprehensive examination of inter-individual variation and intra-individual temporal variation for commonly used measures of intrinsic brain function. We found that definitionally distinct R-fMRI indices of intrinsic brain function tend to exhibit a relatively high degree of co-variation within and between individuals. When taken as a measure of intrinsic brain function, inter-individual differences in concordance for R-fMRI indices appeared to be stable, and negatively related to age (i.e., functional concordance among indices decreased with age). In attempting to understand the functional significance of concordance, we noted that higher concordance was generally associated with higher strengths of the various R-fMRI indices. Finally, temporal dynamic analyses revealed that high concordance states are characterized by increased within- and between-network functional connectivity, suggesting more general variations in network segregation and integration.

Our findings that different R-fMRI indices exhibit commonalities in their patterns of variation across time and individuals have multiple implications. From a methodological perspective, functional concordance is not designed to replace the existing metrics, but quantifies the consistence of a set of commonly used R-fMRI metrics (i.e., fALFF, ReHo, VMHC, DC and GSCorr), which are often treated as if independent, when they are not. The concordance measure allows us to capture consistencies among these base metrics. Thus, at a minimum, studies using multiple metrics would benefit from its inclusion; it can be used to guide the selection of individual index and allows comparing and integrating findings across studies using different indices. The present concordance results may also dissuade investigators from choosing one metric or another across studies, without consideration of their larger context. Furthermore, the concordance index, as a summary metric, can be useful in and of itself. We demonstrated that functional concordance could capture phenotypic traits: it is negatively correlated with age. It also has the potential to be extended to neuropsychiatric disorders.

Beyond insights into study or analytic design, we believe the findings of the present work have potentially profound implications for our understanding of intrinsic brain function. In particular, we found that as concordance among measures varied over time, the degree of functional integration in the brain indexed by the various indices changed as well. This is analogous to the high efficiency states and low efficiency states defined by Zalesky et al. (Zalesky et al., 2014). For periods of high concordance, we found that notably different indices of connectivity (i.e., ReHo, DC, GSCorr, VMHC) and oscillatory power all showed maximal values, suggesting heightened functional integration. Consistent with this notion, connectivity both within and between large-scale functional networks was greater in these states. In contrast, low concordance periods were generally characterized by lower R-fMRI indices and decreased connectivity within and between networks.

Examination of the correlations between the concordance among R-fMRI indices and physiological signals (e.g., cardiac, respiratory) failed to find any meaningful relationships, increasing suspicions of a potential neural etiology for the phenomenon of high concordance. Prior findings suggested that widespread fluctuations in fMRI signals (i.e., global signal) are correlated with spontaneous fluctuations detectable in the electrophysiological local field potentials (Scholvinck et al., 2010). Additionally, dynamic changes in functional connectivity were associated with variations in the amplitude of alpha and theta oscillations measured by electroencephalography (Chang et al., 2013), and are linked to wakefulness (Tagliazucchi and Laufs, 2014). Consistent with these prior works, our findings emphasize the potential value of examining global brain connectivity properties and draw attention to spontaneous fluctuations in the connectedness of brain regions – both within and between functional systems, as well as locally (i.e., ReHo) and globally (i.e., DC, GSCorr). Additionally, our work suggests that these fluctuations in connectivity across brain regions are linked to fluctuations in the prominence of low frequency fluctuations.

As was expected a priori, the findings of the present work verified that functional concordance was much higher in grey matter than white matter or CSF, where little meaningful signal is expected. A key question is what drives differences in functional concordance among individuals. The first possibility is that morphology may be a driving factor. Across individuals, we found progressive age-related decreases across the lifespan (ages: 8-86). During development, reductions in gray matter volume proportion are prominent from childhood to adolescents (Sowell et al., 2002). A rich literature has demonstrated aging-related gray matter loss from adulthood to old age (e.g. Bartzokis et al., 2001; Jernigan et al., 2001). In a lifespan study (ages from 7 to 87 years) (Sowell et al., 2003), gray matter reduction continued across the whole lifespan – a phenomenon we confirmed in our supplementary analyses (i.e., gray matter density negatively correlated with age across the lifespan: r =-0.70, Figure S4). However, controlling for gray matter density did not account for the changes we observed, suggesting a more complex phenomenon. A second possibility is vascular function. Cerebrovascular reactivity measured by CO_2_ inhalation or breath holding declines with age from teenagers to adults (Leung et al., 2016) and across the adult lifespan (Ito et al., 2002; Liu et al., 2013; Lu et al., 2011), but increases from childhood to mid-teens (Leung et al., 2016). Cerebral blood flow is higher in children compared to adults (Leung et al., 2016; Moses et al., 2013; Moses et al., 2014), and decreases in normal aging (Chen et al., 2011; Lu et al., 2011). Reduced blood supply, together with the decreased capacity for vasodilatation, could produce changes in neurovascular coupling that could contribute to the decline in functional concordance over age. Further investigation will be required to investigate the contributions of such phenomena.

The present study revealed a temporal dynamic pattern of regional and global aspects of intrinsic brain activities. This dynamic nature is consistent with previous reports that focused on dynamic functional connectivity patterns (Chang and Glover, 2010; Sakoglu et al., 2010), based on which intrinsic brain activities could be decoded into several states (Allen et al., 2014; Leonardi et al., 2013); see reviews in (Calhoun et al., 2014; Hutchison et al., 2013; Tagliazucchi and Laufs, 2015). Interestingly, high-level cognitive regions, including the default mode network and dorsolateral prefrontal cortex, demonstrated that temporal variations were lower in regional activity measures (i.e., ALFF and fALFF), but higher in functional connectivity indices (i.e., ReHo and DC). These regions are cortical hubs, and thus are expected to exhibit greater stability. This is supported by the high test-retest reliability of these regions (Zuo and Xing, 2014), and further supported by their lower temporal variability in regional strength. However, these regions are highly connected to other brain regions to process multimodal information (Buckner et al., 2009); thus the connections should be flexible. This is supported by high temporal variability in functional synchrony measures in the current study and a prior study (Zhang et al., 2016), as well as by findings suggesting that these regions consistently form dynamic functional connections (Zalesky et al., 2014). In contrast, the lower-level primary regions showed higher temporal variability in both regional strength and functional synchrony (high CV in ALFF, fALFF, ReHo and DC). The actual regional activity (as indexed by ALFF) in primary motor regions was relatively lower than high-level cognitive regions such as dorsolateral prefrontal cortex. This indicates lower-level regions maintain less active but more flexible states during the resting-state. Finally, the high CV for fALFF observed within the subcortical and limbic areas is of particular interest, as these areas tend to be characterized by lower test-retest reliability. This may be reflective of lower signal to noise ratio, though it can also suggest greater context dependencies for connectivity properties of these areas (Craddock et al., 2013b; Mennes et al., 2013). Of note, these areas also showed greater prominence of age-related reductions, as well as modest increases across concordance states – again pointing away from a simple scanner-related SNR explanation.

Several limitations in the present work merit consideration. First, the five R-fMRI indices we selected were considered to reflect different functional aspects of intrinsic brain activity. Other R-fMRI indices (e.g., various seed-based correlation maps, ICA components, Hurst exponent) not examined here may facilitate a deeper understanding of the physiological processes underlying intrinsic brain activity. Second, we found ensemble changes of intrinsic brain activity which could be divided into high-integration and low-integration states. As simultaneous electrophysiological recordings were not available for these datasets, we can only speculate how neuronal ensemble firing might relate to these fluctuations in functional concordance. Third, while we found that functional concordance is related to age in healthy participants, its generalizability to clinical populations remains unknown. Future studies should apply these methods to neuropsychiatric disorders to test their sensitivity and specificity in brain disorders. Finally, we note that challenges associated with establishing the potential contributions of factors such as scanner drift or the adaptation of participants to the scanning environment. In the present study, we observed that across participants, concordance tended to increase slightly over time. Although this was minimally detectable at the individual level and unlikely to contribute to our findings, the etiology of such changes are not clear and worthy of future study.

In sum, the present work comprehensively examined of the functional concordance of intrinsic brain indices from inter-individual variation and temporal dynamics perspectives, provided new insights to examine the differencing states within and between individuals. Our work suggests global fluctuations exist in the brain, and their spontaneous variations over time result in fluctuations in the connectedness of brain regions.

## ACKNOWLEDGEMENTS

The authors appreciate the editorial assistance and support of Dr. Francisco X. Castellanos. This work was supported by the National Key R&D Program of China (2017YFC1309902 to CGY), National Basic Research (973) Program (2015CB351702 to XNZ), the Natural Science Foundation of China (81671774 and 81630031 to CGY, 81471740, 81220108014 to XNZ), the Hundred Talents Program of the Chinese Academy of Sciences (Y5CX072006 to CGY), Beijing Municipal Science & Technology Commission (Z161100000216152 to CGY), the National Institutes of Health (U01MH099059 to MPM), the Child Mind Institute (1FDN2012-1 to MPM) and gifts to the Child Mind Institute (MPM) from Phyllis Green, Randolph Cowen, and Joseph P. Healey.

---

**Figure S1.** Differences in magnitude of R-fMRI indices between high and low concordance participants or time windows (between the upper tercile and the lower tercile). The figure layout is the same as Figure 6. The differences is that the computation was based on upper tercile and the lower tercile instead of upper and lower 20%.

**Figure S2.** Effect size (Cohen’s f^2^) of differences in magnitude of R-fMRI indices between high and low concordance participants or time windows (between the upper 20% and the lower 20%). The figure layout is similar to Figure 6, while putting Cohen’s f^2^ maps instead of T maps.

**Figure S3.** The impact of scanning temporal order on functional concordance based on the phase randomized surrogate data. The figure layout is the same as Figure 9. The difference is that the computation is based on surrogate data instead of real data. There were no prominent linear trend appear on all R-fMRI indices for the surrogate data.

**Figure S4.** The correlation between global gray matter density (averaged across voxels within the grey matter mask) and age across the lifespan.

1 The group mask was generated using DPABI’s Quality Control module. Specifically, for each participant, DPABI generates a coverage mask based on the mean EPI image which was then registered into MNI space. The voxels present in at least 90% of participants were included in the group mask.

2 Window sizes of 64 TRs and 128 TRs were also tested, resulting in similar results.

3 A group GM mask was used to avoid including ventricular and white matter signals. The GM mask was generated by averaging the GM density maps across participants, and then thresholding at 0.33 and making an intersection with the group mask (90% on EPI converge).

4 The smoothing in preprocessing was only performed for VMHC (4.5 mm FWHM) but not other R-fMRI indices, because VMHC needs a good correspondence between symmetric voxels, which requires smoothing (in addition to being registered to a symmetric template). In group analyses, to improve the correspondence of a voxel across participants (e.g., minimize the variation induced by registration error), smoothing was performed on the derivatives right before group analysis. Here, the static R-fMRI indices were smoothed (4.5mm FWHM) and then Kendall’s W of the five static R-fMRI indices across participants was calculated.

5 Kendall’s W of the five dynamic R-fMRI indices across time was calculated on unsmoothed R-fMRI index maps and then the Kendall’s W maps were smoothed (4.5mm FWHM). Then one-sample t-test was performed on these smoothed concordance maps after Z-standardization (subtracting the global mean and dividing by the standard deviation).

6 To ensure that our results were not biased by our decision to sort the data into quintiles, we repeated this analysis by dividing the participants into terciles and comparing participants with the highest 1/3 volume-wise static concordance with those with lowest 1/3; as demonstrated in Figure S1, we found a highly similar pattern of results.

